# *Yersinia pseudotuberculosis* doxycycline tolerance strategies include modulating expression of genes involved in cell permeability and tRNA modifications

**DOI:** 10.1101/2021.11.01.466809

**Authors:** Hector S. Alvarez-Manzo, Robert K. Davidson, Jasper Van Cauwelaert de Wyels, Katherine L. Cotten, Benjamin Nguyen, Zeyu Zhu, Jon Anthony, Tim van Opijnen, Kimberly M. Davis

## Abstract

Antibiotic tolerance is typically associated with a phenotypic change within a bacterial population, resulting in a transient decrease in antibiotic susceptibility that can contribute to treatment failure and recurrent infections. Although tolerant cells may emerge prior to treatment, the stress of prolonged antibiotic exposure can also promote tolerance. Here, we sought to determine how *Yersinia pseudotuberculosis* responds to doxycycline exposure, to then verify if these gene expression changes could promote doxycycline tolerance in culture and in our mouse model of infection. Only four genes were differentially regulated in response to a physiologically-relevant dose of doxycycline: *osmB* and *ompF* were upregulated*, tusB* and *cnfy* were downregulated; differential expression also occurred during doxycycline treatment in the mouse. *ompF, tusB* and *cnfy* were also differentially regulated in response to chloramphenicol, indicating these could be general responses to ribosomal inhibition. *cnfy* has previously been associated with persistence and was not a major focus here. We found deletion of the OmpF porin resulted in increased antibiotic accumulation, suggesting expression may promote diffusion of doxycycline out of the cell, while OsmB lipoprotein had a minor impact on antibiotic permeability. Overexpression of *tusB* significantly impaired bacterial survival in culture and in the mouse, suggesting that tRNA modification by *tusB*, and the resulting impacts on translational machinery, may play an important role in promoting tolerance. We believe this is the first observation of bactericidal activity of doxycycline, which was revealed by reversing *tusB* downregulation.

## Introduction

Antibiotics have saved countless lives, however since the introduction of penicillin as a clinical treatment in the 1940s, researchers and clinicians observed that bacterial pathogens may have inherent resistance or decreased susceptibility to antibiotics ^1, 2^. This can occur through many different molecular mechanisms, which are broadly grouped into antibiotic resistance and antibiotic tolerance mechanisms. Antibiotic resistance is typically associated with a genetic change within a bacterial population, which allows cells to not only survive, but continue to replicate, in the presence of inhibitory concentrations of antibiotics ^3, 4^. In contrast, antibiotic tolerance is typically associated with transient, phenotypic changes, though genetic changes can also contribute to tolerance ^3–6^. During antibiotic treatment, tolerant cells survive in the presence of inhibitory concentrations of drug, but do not divide until drug concentrations decrease to subinhibitory levels ^3, 4, 7^. Antibiotic tolerance can be caused by environmental stressors imparted on the bacterial population prior to antibiotic exposure, but tolerance can also emerge in response to exposure to the drug itself ^8–11^. Generally, antibiotic tolerance is associated with transient slowed growth ^8, 12, 13^, which renders antibiotic action less effective. Although resistance is more heavily studied, tolerance also plays a major role in antibiotic treatment failure, and it is critical to better understand tolerance processes to develop novel therapeutic options that more effectively eliminate bacterial infection.

Antibiotics generally fall into two categories: bactericidal and bacteriostatic drugs; where bactericidal drugs directly kill bacteria and bacteriostatic drugs arrest bacterial growth. Bacteriostatic drugs then rely on the host immune system to clear growth-arrested cells to resolve infection. While definitions of antibiotic resistance and tolerance readily apply to bactericidal drugs, bacteriostatic drugs inherently arrest growth of bacterial population, and so all bacterial populations would be considered ‘tolerant’ to bacteriostatic drugs. This definition of tolerance then only applies to bacteriostatic drugs that have bactericidal activity under a particular condition ^14–16^. This is an important distinction for this study, which focuses on doxycycline treatment. Doxycycline was chosen for these studies because it is used to treat *Yersinia* infections, we have an established mouse model to study its efficacy, and fluorescent reporters for tracking doxycycline exposure ^17–22^. Doxycycline is a bacteriostatic antibiotic of the tetracycline class that binds reversibly to 16S rRNA ^23–25^. Two different mechanisms are known to promote doxycycline resistance: expression of efflux pumps that lower intracellular drug concentration, and production of proteins that protect ribosomes from doxycycline binding ^23, 25^. Doxycycline tolerance has never been explored because bactericidal activity has not been described. However, in this study, we have observed bactericidal activity of doxycycline during treatment of a mutant strain, which suggests that our wild-type (WT) strain can be defined as doxycycline-tolerant. We believe this may be the first description of bactericidal activity of doxycycline, and could be related to altered binding of doxycycline to ribosomes in this strain.

Bacterial infections are particularly difficult to treat when bacteria spread to the bloodstream and colonize deep tissue sites, such as the spleen. It can be difficult to deliver inhibitory concentrations of antibiotics to deep tissues, and it can be challenging to eliminate the entire bacterial population ^17, 26, 27^. Subsets of bacteria may survive due to exposure to subinhibitory concentrations of antibiotics, and subsets of bacteria may express sets of genes that render them less susceptible to antibiotics. When a subpopulation of bacteria emerges with increased antibiotic tolerance, this is termed antibiotic persistence, and the subpopulation is comprised of persister cells ^4, 7^. *Yersinia pseudotuberculosis* is known to cause long-term infections, associated with both intestinal and deep tissue sites, but it remained unclear if antibiotic persistence was underlying these clinical observations ^28–30^. Our recent study showed that approximately 10% of the *Y. pseudotuberculosis* population replicating in the spleen will survive a single inhibitory dose of doxycycline (Dox) ^17^. Dox was chosen for these experiments because it should be an effective treatment for *Yersinia* infection ^18, 22, 31^. However, we found this bacterial subpopulation persisted in host tissues ∼48 hours until the Dox level decreased, and then resumed growth, which defines these cells as an antibiotic-tolerant subpopulation of persister cells ^3, 32^. Stressed bacterial cells expressing the nitric oxide detoxifying gene, *hmp*, were enriched in the surviving cells, indicating NO stress may cause a change in bacteria that promotes survival ^17^. However, only ∼40% of surviving cells were Hmp^+^, suggesting additional pathways also contribute to survival. Exposure to subinhibitory doses of antibiotics can also promote survival by altering gene expression in the surviving cells ^8, 9^. Since this surviving bacterial population is exposed to inhibitory concentrations of Dox for ∼48 hours prior to resuming growth, we hypothesized that phenotypic changes in response to Dox may also contribute to survival.

In this study we sought to determine if differential expression of specific genes alters the doxycycline susceptibility of *Y. pseudotuberculosis*. We performed RNA-seq to identify genes with altered expression during exposure to Dox, and constructed deletion and overexpression strains to determine the role of these gene products in *Y. pseudotuberculosis* doxycycline susceptibility. We show here that loss of expression of two outer membrane proteins, OmpF and OsmB, altered doxycycline accumulation, although this had minimal impact on bacterial survival. We also show that heightened expression of *tusB,* whose gene product is predicted to increase translational fidelity by stabilizing the wobble position of tRNAs ^33, 34^, significantly increases Dox sensitivity.

## Results

### *ompF, osmB, CNFy,* and *tusB* are differentially regulated in response to doxycycline

The phenotypic changes that promote antibiotic tolerance can occur prior to antibiotic treatment or in response to antibiotic treatment. Our recent study showed there is a subpopulation of *Y. pseudotuberculosis* that survives doxycycline (Dox) treatment in the mouse spleen, and host-derived nitric oxide initially promotes survival of stressed cells ^17^. This tolerant subpopulation persists within host tissues for 48 hours before Dox concentrations wane and the bacterial population resumes growth. We hypothesized there may be phenotypic changes that occur in response to Dox, which promote bacterial survival in the hours after treatment.

Using a fluorescent transcriptional reporter, we have approximated the concentrations of Dox in the spleen at 0.1µg/ml-1µg/ml during this period of growth inhibition following a single injection of 40mg/kg Dox ^17^. To determine if there are specific pathways upregulated or downregulated during exposure to Dox, we initially chose to expose bacteria to 0.1µg/ml Dox. All bacterial cells would be exposed to at least this concentration within mouse tissues, and this concentration was sufficient for maximal de-repression of the *tet* operon ^17^, so we expected to see many transcriptional changes. We initially tested several different genes in known stress response pathways (ROS, RNS, general stress) to determine if these were differentially regulated in Dox-exposed *Y. pseudotuberculosis* in culture, but did not find significant changes **(Supplemental Figure 1)**. We then took a global approach and utilized RNA-Seq to determine if there are specific pathways that would be up- or downregulated during exposure to 0.1µg/ml Dox. We exposed broth-grown bacteria to Dox and isolated bacterial RNA after 2h (hours, h) and 4h Dox exposure; treated and untreated cultures were grown in parallel. Samples were prepared for sequencing using a RNAtag-seq approach ^35^ and aligned to the *Y. pseudotuberculosis* genome. Although this concentration of Dox strongly modulates the TetR *tet* repressor ^17^, very few transcriptional changes occurred during antibiotic exposure *in vitro* (**Figure 1A**). We identified 8 genes that were differentially regulated after 2h Dox treatment (**Figure 1B**), and there were no significant changes after 4h treatment, suggesting that transcriptionally, cells had returned to baseline. Of the 8 genes, only two genes were upregulated ∼2 fold *(osmB, ompF*), and two genes were downregulated ∼2 fold *(tusB, cnfy*), and so these 4 genes were selected for downstream analyses (highlighted in red, **Figure 1B**). We confirmed these 4 genes were differentially regulated after 2h Dox exposure by prepping additional bacterial samples for qRT-PCR, and also showed that gene expression changes were no longer observed after 4h Dox exposure (**Figure 1C**). These results were surprising; based on previous studies we had expected to see increased expression of efflux pumps and decreased expression of porins ^36–38^, instead we saw upregulation of a porin (*ompF*) and no change in efflux pump expression.

**Figure 1:**
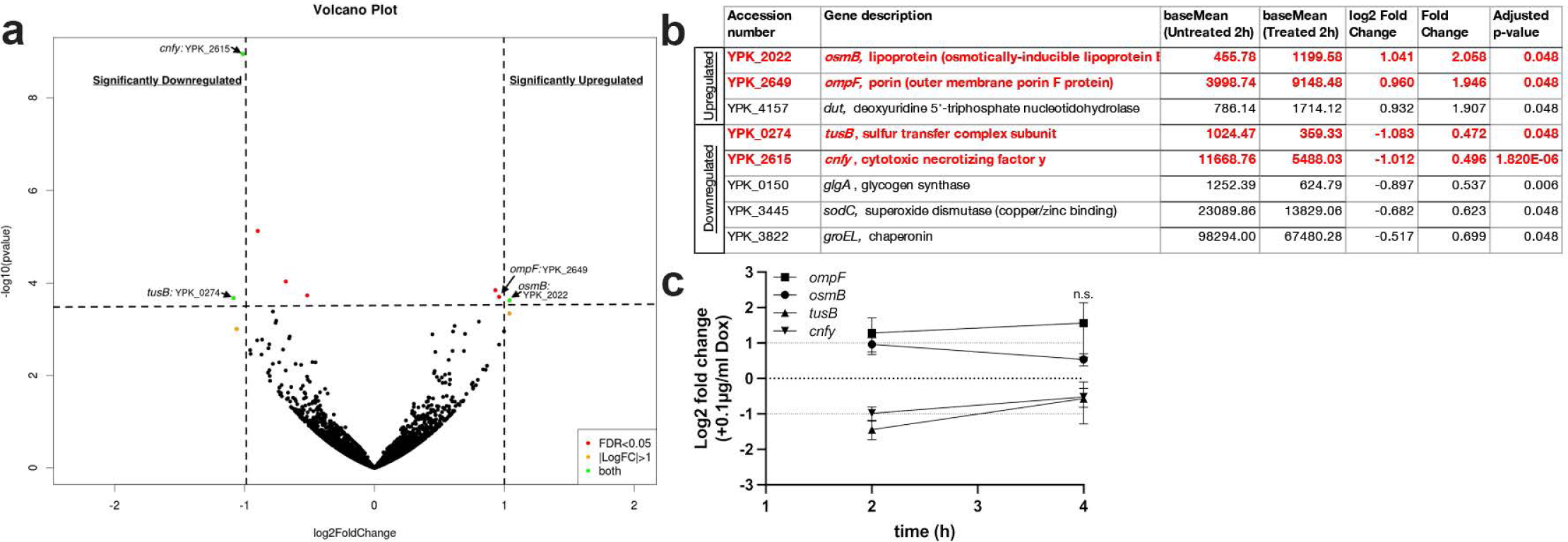
*ompF, osmB, CNFy,* and *tusB* are differentially regulated in response to doxycycline. Cultures of WT *Y. pseudotuberculosis* were incubated in the presence or absence of 0.1µg/ml Dox and bacterial transcript levels were compared by RNA-seq. Significant changes in transcript levels were detected at 2h post-treatment by DESeq2 analysis. **(A)** Volcano plot of aligned genes, each dot represents one gene. Horizontal dotted line: adjusted p value of 0.05, genes above this line are considered significantly altered. X-axis: log2 fold change in transcript levels. Vertical lines mark 2-fold changes, genes outside these lines are differentially regulated more than 2-fold. Hits of interest are highlighted. **(B)** Table of significant hits, 2h post-treatment. Hits of interest (differentially regulated ∼2-fold) are highlighted in red. Base mean values represent 4 biological replicates. Log2 fold change and fold change are shown with adjusted p-value (adjusted based on gene length). **(C)** qRT-PCR validation of RNA-seq results. Cultures were prepared as described above, and bacterial transcripts were detected by qRT-PCR. Log2 fold change values were calculated relative to untreated cells, horizontal dotted lines depict average values for untreated cells (0log2), and 2-fold changes. Data represents four biological replicates. Statistics: **(C)** Two-way ANOVA with Tukey’s multiple comparison test, comparisons made relative to untreated cells. n.s.: not significant.

### *ompF, osmB, CNFy,* and *tusB* are differentially regulated in response to doxycycline *in vivo*

Many studies have characterized how bacteria respond to antibiotics during growth in bacteriological media, however, few studies have identified changes that occur within the bacterial population during antibiotic treatment within host tissues. To determine if the gene expression changes we observed in culture are relevant in our mouse model of systemic infection, we infected C57BL/6 mice intravenously with WT *Y. pseudotuberculosis*, allowed infection to proceed 48h, and delivered a single injection of 40mg/kg Dox intraperitoneally ^17^. Spleens were isolated at 48h (0h, time of Dox administration), or at 2h or 4h post-treatment.

RNA was stabilized, isolated, mammalian mRNAs, tRNAs, and rRNAs were depleted to enrich for bacterial RNA, and bacterial transcript levels were detected by qRT-PCR. All 4 genes were differentially regulated in the host environment and the fold change in transcript levels was more dramatic within host tissues than seen with 0.1µg/ml Dox exposure (**Figure 2A**). *osmB* and *cnfy* had significant changes in transcript levels after 4h treatment, compared to the time of Dox administration (0h) (**Figure 2A**). For *tusB,* transcripts were significantly lower at 2h post-treatment, but not at 4h. Although there was high upregulation of *ompF* transcript levels, this was not statistically higher than baseline (0h), possibly due to some variability across samples.

**Figure 2:**
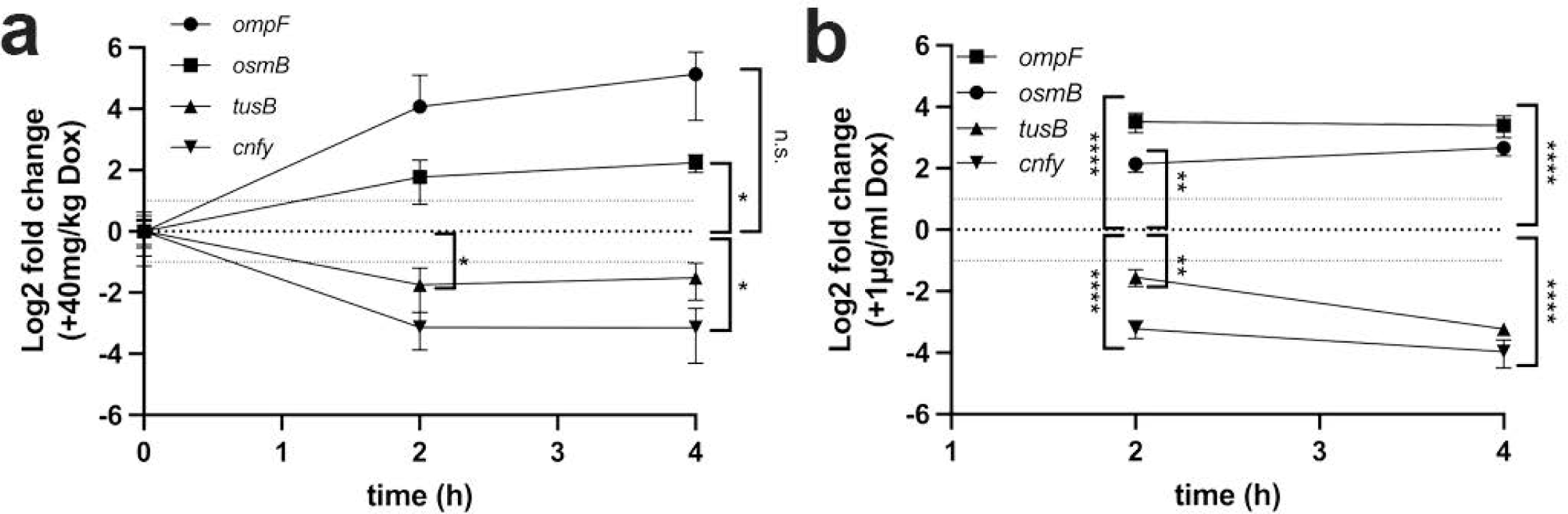
*ompF, osmB, CNFy,* and *tusB* are differentially regulated in response to doxycycline *in vivo*. **(A)** C57BL/6 mice were infected with WT *Y. pseudotuberculosis*, infection proceeded for 48h (hours, h), and mice were injected intraperitoneally with a single dose of 40mg/kg Dox. At the indicated timepoints spleens were harvested, and samples were enriched for bacterial RNA prior to qRT-PCR. Time: h post-treatment, 0h represents 48h post-inoculation (p.i.), the time of treatment. Log2 fold change is relative to 0h. Horizontal dotted lines depict average values for 0h (0log2), and 2-fold changes. Data represents 4 biological replicates/group. **(B)** Cultures of WT *Y. pseudotuberculosis* were incubated for 4h in the presence or absence of 1µg/ml Dox, bacterial RNA was isolated and transcripts were detected by qRT-PCR. Log2 fold change values were calculated relative to untreated cells, horizontal dotted lines depict average values for untreated cells (0log2), and 2-fold changes. Data represents four biological replicates. Statistics: (**A)** Two-way ANOVA with Bonferroni’s multiple comparison test, comparisons made relative to 48h p.i./time 0h timepoint. **(B)** Two-way ANOVA with Tukey’s multiple comparison test, comparisons made relative to untreated cells using relative expression values. ****p<.0001, ***p<.001, **p<.01, *p<.05, n.s.: not significant.

Based on our previous study, we believe bacteria are exposed to 0.1-1µg/ml Dox within the mouse spleen in this treatment model ^17^. To determine if these 4 genes are also differentially regulated in response to 1µg/ml Dox in culture, we performed additional *in vitro* experiments and isolated RNA following 2h and 4h exposure to 1µg/ml Dox to quantify transcript levels by qRT-PCR. All 4 genes were differentially regulated at 2h post-treatment relative to untreated cells, and in contrast with 0.1µg/ml treatment, this differential regulation was maintained 4h post-treatment (**Figure 2B**). The magnitude of change in transcript levels was also much higher with 1µg/ml compared to 0.1µg/ml Dox, and was between an 8-16-fold change at 4h post-treatment with 1µg/ml Dox (**Figure 2B**), compared to a maximum 2-fold change with 0.1µg/ml Dox (**Figure 1B**). The change in transcript levels within the mouse spleen fell in the middle of these two treatment condition values (**Figure 2A**), consistent with a range of exposure between 0.1-1µg/ml Dox in the spleen.

OmpF is a well described bacterial porin, which is localized in the outer membrane of Gram-negative bacteria, and plays a major role in antibiotic diffusion into the cell ^23, 36, 38^. Interestingly, mutations in *ompF* that either reduce functionality or downregulate expression of the gene have been strongly associated with resistance to β-lactams ^39, 40^. Our results instead suggest that upregulation of *ompF* could be protective during antibiotic exposure in *Yersinia*. *osmB* encodes for an outer membrane lipoprotein, and its expression is regulated by changes in osmolarity in *E. coli* ^41, 42^. Expression of *osmB* is also regulated by the alternative sigma factor, RpoS, and increases in stationary phase *E. coli* ^43, 44^, but *osmB* has not been previously associated with antibiotic susceptibility. However, increased expression of *osmB* in stationary phase could suggest some association of this gene product with slowed growth. TusB functions with TusC and TusD in the sulfur-relay system, which has been associated with translational fidelity, through proper incorporation of glutamic acid, glutamine and lysine into growing polypeptide chains by stabilizing the wobble position of tRNAs ^33^. TusB has not been previously associated with antibiotic susceptibility or tolerance. CNFy has been previously associated with persistence in *Y. pseudotuberculosis* ^28^, and for that reason, it will not be a focus in the preceding figures.

### *ompF* mutants have slightly impaired growth in the presence of doxycycline

The antibiotic susceptibility of a bacterial strain is assessed by several well-defined measurements. These include the minimum inhibitory concentration (MIC): the concentration required for complete bacterial growth inhibition during culture; the minimum bactericidal concentration (MBC): the concentration required for a certain amount of bacterial killing over a defined time period, usually 90% killing or higher; and the more recently described minimum duration for killing (MDK): the amount of time required to kill a bacterial population during exposure to inhibitory levels of drug, typically used to assess the antibiotic tolerance of a bacterial strain ^4, 5, 45–47^. Using this framework, we set-up three sets of *in vitro* assays to assess the antibiotic susceptibility of our WT and mutant strains.

To determine if increased expression of *osmB* or *ompF* promotes antibiotic tolerance, we generated single deletion mutants that lack either gene product (Δ*osmB,* Δ*ompF*), and a double deletion mutant strain that lacks both genes (Δ*osmB* Δ*ompF*). We first wanted to test the antibiotic susceptibility of the mutant strains by performing growth curves to determine if the MIC was significantly altered by a loss of these gene products. The MIC should only be altered by the presence or absence of genes required for antibiotic resistance, while gene products contributing to tolerance should not impact the MIC of the strain ^4^. Since the WT MIC is 1µg/ml ^17^, and we would expect mutants to show no change or increased sensitivity, the highest dose used here was 1µg/ml. Under untreated conditions, we found the Δ*osmB* Δ*ompF* strain was already growing slightly slower than the WT strain (**Figure 3A**). This strain was also growing slower under treated conditions (0.01-1µg/ml) (**Figure 3B-D**), suggesting growth rates may be slowed in this strain. We found very little impact on the growth of the Δ*osmB* strain, but interestingly, the Δ*ompF* strain did appear slightly more sensitive to Dox when compared to the WT strain (0.01-0.1µg/ml) (**Figure 3B, 3C**). During exposure to inhibitory concentrations (1µg/ml) of doxycycline, the absorbance of the Δ*ompF* strain was higher than WT, albeit significantly inhibited. Given the growth inhibition of Δ*ompF*, we would have expected the Δ*osmB* Δ*ompF* strain to have similar growth inhibition, however, these results were the first indication that the *osmB* deletion may partially rescue the Δ*ompF* phenotype. These slight changes in growth of the mutant strains were not sufficient to alter their MICs.

**Figure 3:**
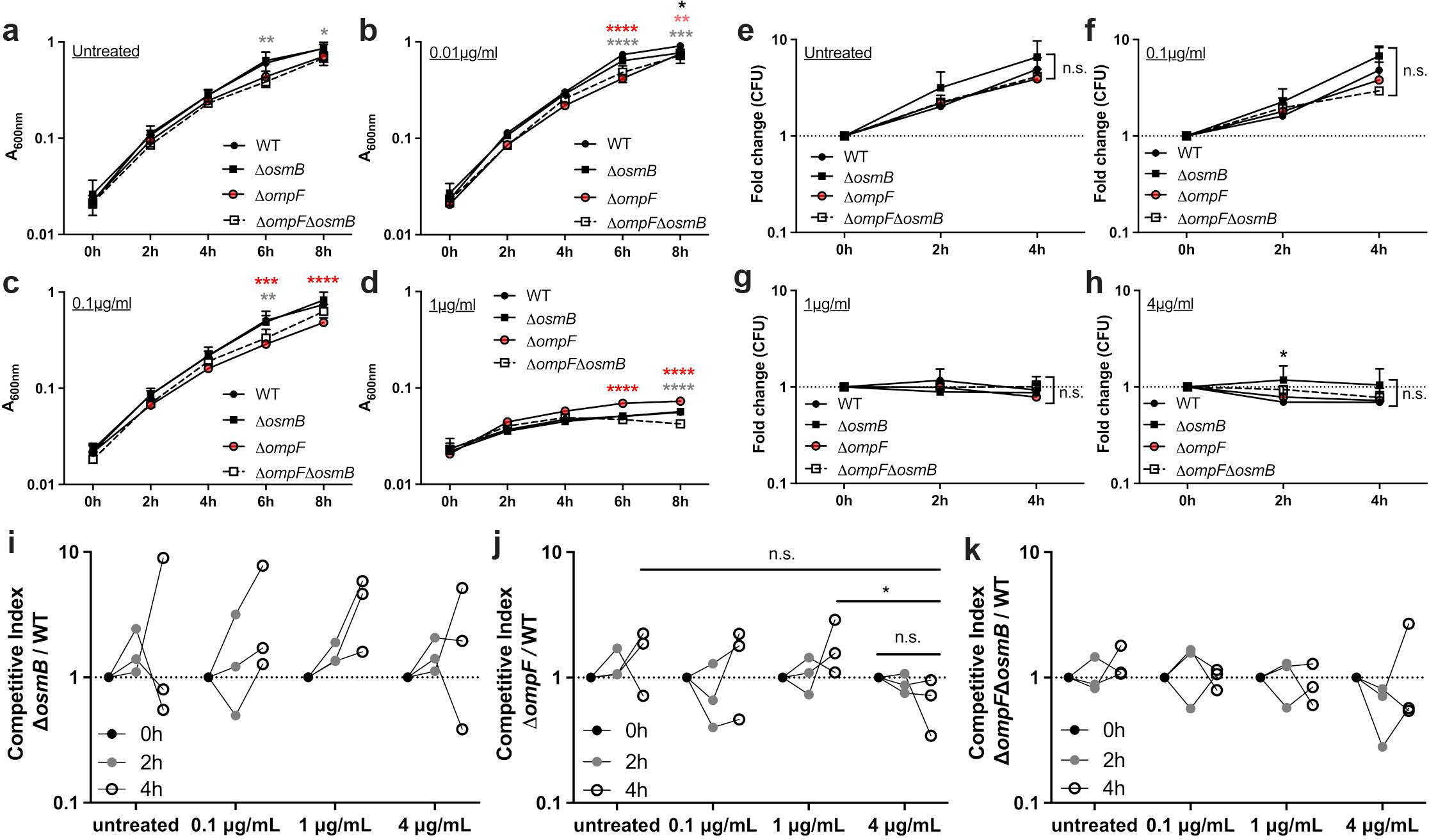
*ompF* mutants have slightly impaired growth in the presence of doxycycline. Exponential phase cultures of the WT and mutant strains (Δ*osmB,* Δ*ompF,* Δ*osmB* Δ*ompF*) were treated with the indicated concentrations of Dox to assess **(A-D)** growth inhibition, **(E-H)** changes in viable counts, and **(I-K)** competitive survival. **(A-D)** Strains were incubated with the indicated concentrations of Dox and growth inhibition was assessed based on absorbance (A_600nm_) at the indicated timepoints (time: hours, h). Data represents the mean and standard deviation of three biological replicates. **(E-H)** Strains were incubated with the indicated concentrations of Dox and viable counts were determined by quantifying CFUs. Fold change in CFUs is shown relative to time 0h. Dotted line: a value of 1, no change relative to 0h. Data represents the mean and standard deviation of three biological replicates. **(I-K)** Competitive survival in the presence of the indicated concentrations of Dox; single and double mutants were tested alongside the WT strain. Competitive index (CI): CFUs of the mutant/WT divided by the ratio of mutant/WT in the culture at time 0h. Values above 1 indicate the mutant preferentially survives, values less than 1 indicate the WT preferentially survives. Dotted line: value of 1, equal fitness. Dots: biological replicates, lines connect biological replicates sampled across the timepoints, three biological replicates shown. Statistics: Two-way ANOVA with Tukey’s multiple comparison test, **(A-H)**: comparisons made relative to the WT strain; **(J)**: comparisons made between 4h treatment CI values. ****p<.0001, ***p<.001, **p<.01, *p<.05, n.s.: not significant.

We then tested the viability of the WT and mutant strains during exposure to heightened, inhibitory levels of doxycycline (0.1-4µg/ml). The highest concentration used here was 4µg/ml, since this is the approximate peak concentration in serum in the mouse model ^19, 48, 49^. We exposed exponential phase cells to doxycycline for 4h, and plated cells at the point of adding doxycycline (0h), 2h after exposure, and 4h after exposure to determine whether bacteria continued to grow, were growth inhibited, or lost viability during the exposure. Each strain was grown in a separate test tube with rotation, and we calculated the fold change in CFUs relative to the 0h timepoint. Strains continued to grow under untreated conditions and during exposure to 0.1µg/ml doxycycline **(Figure 3E, 3F**), while all strains were growth-inhibited during exposure to 1-4µg/ml doxycycline (**Figure 3G, 3H**). There were not significant differences in the viability of the strains, with the exception of the Δ*osmB* strain, which had significantly more CFUs than WT at 2h treatment with 4µg/ml. These data also suggest there is not a detectable difference in the MIC of the mutant strains relative to the WT, and again indicate the MIC for all strains is 1µg/ml. Because bacterial killing was not observed, the MBC or MDK cannot be determined based on these results.

We then tested *in vitro* doxycycline sensitivity in one additional assay, designed to test competitive survival of two co-cultured strains. The WT strain was transformed with a plasmid that contains a chloramphenicol resistance cassette and also constitutive GFP signal ^50, 51^. Mutants lacked an antibiotic resistance cassette, allowing us to identify the abundance of each strain by plating on selective and non-selective media. Strains were grown to exponential phase then added to individual wells of a 96-well plate. Mutant strains were tested for competitive survival during co-culture with the WT strain, and single strain controls were also included to assess growth and survival of individual strains during doxycycline exposure in the plate, to ensure this mirrored the viability results. Competitive index values were calculated based on the ratio of mutant/WT CFUs at the indicated timepoints after treatment relative to the ratio of mutant/WT CFUs in the well at 0h, when doxycycline was added. Data comparing survival of the WT and Δ*osmB* strain suggested that the Δ*osmB* may have a slight fitness advantage relative to the WT strain, and showed no increased doxycycline sensitivity (**Figure 3I**). Consistent with the slightly impaired growth of the Δ*ompF* strain **(Figure 3B, 3C**), Δ*ompF* showed decreased relative survival during exposure to 4µg/ml doxycycline, suggesting this strain does have slight doxycycline sensitivity (**Figure 3J**). Interestingly, the Δ*osmB* Δ*ompF* strain had very similar survival compared to the WT strain (**Figure 3K**), suggesting this strain does not have doxycycline sensitivity, and again suggesting that the *osmB* deletion may partially rescue the Δ*ompF* phenotype. Collectively, these data suggest that Δ*ompF* strain has increased doxycycline sensitivity, however the *osmB* deletion has no detectable impact on doxycycline sensitivity, and may actually improve the fitness of the strain.

### Δ*osmB* has increased *in vivo* fitness relative to the WT strain, Δ*ompF* does not appear significantly more susceptible to doxycycline *in vivo*

To determine if the Δ*osmB* or Δ*ompF* strains had increased doxycycline sensitivity *in vivo*, we tested their survival in our mouse model of infection during exposure to a single dose of doxycycline. Because the *in vitro* phenotypes were subtle, we decided to test the survival of mutant strains relative to the WT in a co-infection experiment, essentially mirroring the competitive survival experiments we had set-up in culture **(Figure 3I, 3J**). Mutant strains used for infection contained a *yopE::mCherry* construct that would allow us to detect mutant cells with mCherry fluorescence ^51, 52^. The Δ*osmB* Δ*ompF* strain was not tested here since it showed no difference in antibiotic susceptibility relative to the WT strain (**Figure 3K**).

Equal amounts of the WT and mutant strain were mixed, mice were inoculated intravenously, and 48h post-inoculation mice received a single dose of doxycycline or were left untreated. 24h after doxycycline injection (+Dox 24h), spleens were harvested to quantify CFUs and examine microcolony areas by fluorescence microscopy. The WT strain was marked with a constitutive GFP plasmid and chloramphenicol resistance, and the mutants contained *yopE::mCherry* to mark all mutant cells with mCherry signal, as described above. Total CFU/spleen were first quantified to ensure doxycycline treatment significantly decreased bacterial CFUs, then the numbers and microcolony areas of each strain were quantified. Although the doxycycline treatment significantly decreased CFUs (**Figure 4A**), the Δ*osmB* strain outcompeted the WT strain in the presence and absence of doxycycline (**Figure 4B**), again showing the Δ*osmB* strain does not have increased doxycycline sensitivity, and actually has a fitness advantage relative to WT. Consistent with this, the microcolony areas of the Δ*osmB* strain appeared slightly larger than WT, although this was not statistically significant (**Figure 4C**). The number of microcolonies decreased with treatment for both the WT and Δ*osmB* strains, but microcolonies did not significantly decrease in size after treatment for either strain (**Figure 4C**). We were expecting to see decreased microcolony areas after treatment, but that may have been difficult to assess without comparing to the areas at the time of injection (48h) ^17^. We then quantified the CFUs of each strain before and after treatment within the same tissues. There were more Δ*osmB* CFUs prior to and after treatment, and both WT and Δ*osmB* CFUs significantly dropped in response to treatment (**Figure 4D**).

**Figure 4:**
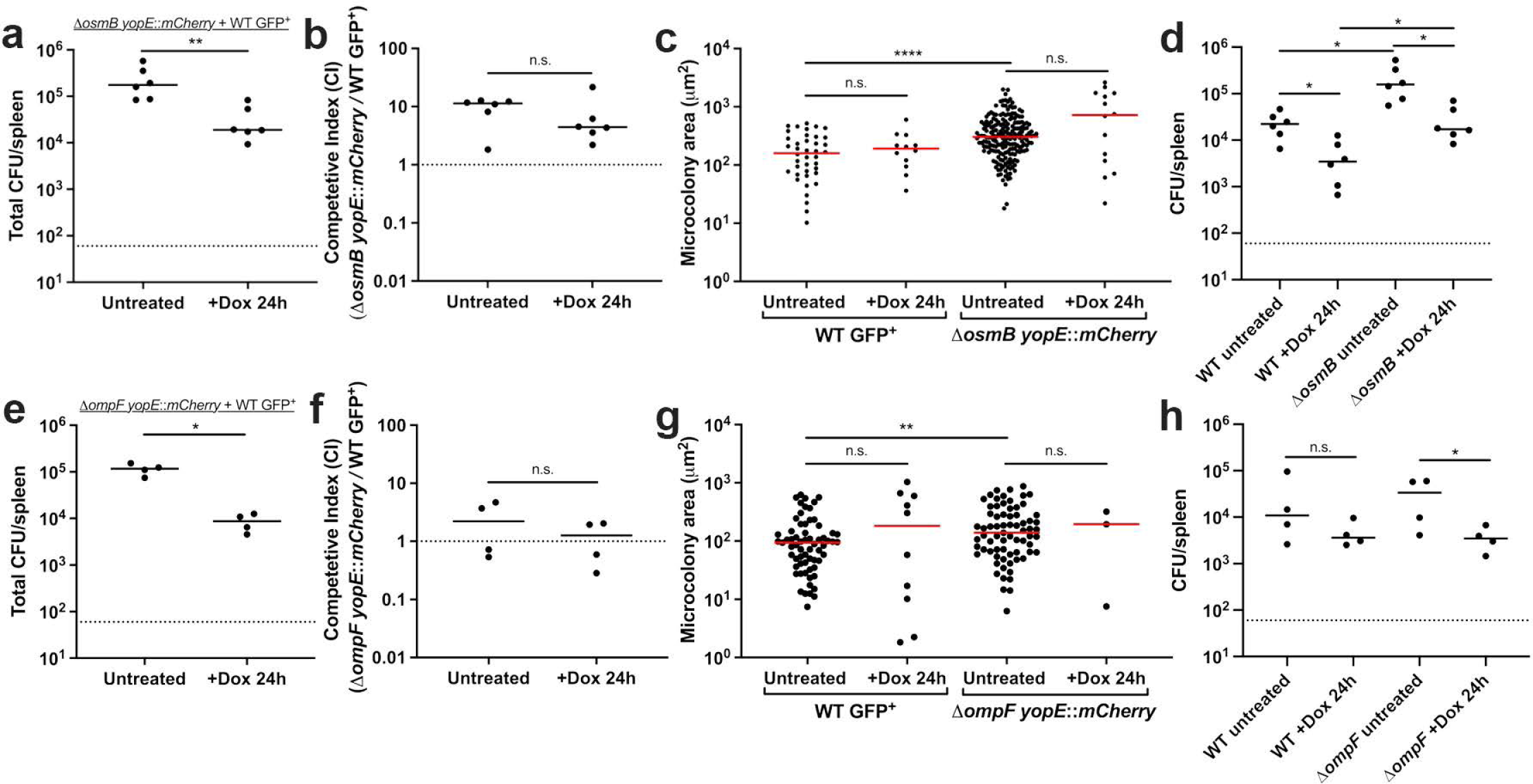
Δ*osmB* has increased *in vivo* fitness relative to the WT strain, Δ*ompF* does not appear significantly more susceptible to doxycycline *in vivo.* C57BL/6 mice were infected with equal numbers of WT and mutant *Y. pseudotuberculosis*, infection proceeded for 48h (hours, h), and mice were injected intraperitoneally with a single dose of Dox or left untreated. Spleens were harvested 24h later to quantify CFUs and microcolony areas. **(A)** Co-infection with Δ*osmB yopE::mCherry* + WT GFP^+^. Total CFU/spleen for treated mice (+Dox 24h) compared to untreated mice at the same timepoint. Dots: individual mice. **(B)** Competitive index: CFUs of the mutant/WT in the spleen divided by the ratio of mutant/WT in the inoculum. Values above 1: mutant is more fit. Dotted line: value of 1, equal fitness. Dots: individual mice. **(C)** Microcolony areas were quantified for individual strains within the harvested spleens (6 untreated mice and 6 treated mice). 20-62 microcolonies were quantified in one section for untreated mice, and 1-13 microcolonies were quantified in one section for treated mice. **(D)** CFU/spleen of each strain in the Δ*osmB* + WT competition experiments. (**E**) Co-infection with Δ*ompF yopE::mCherry* + WT GFP^+^. Total CFU/spleen for treated mice (+Dox 24h) compared to untreated mice at the same timepoint. Dots: individual mice. **(F)** Competitive index: CFUs of the mutant/WT in the spleen divided by the ratio of mutant/WT in the inoculum. Values above 1: mutant is more fit. Dotted line: value of 1, equal fitness. Dots: individual mice. **(G)** Microcolony areas were quantified for individual strains within the harvested spleens (4 untreated mice and 4 treated mice). 14-72 microcolonies were quantified in one section for untreated mice, and 1-7 microcolonies were quantified in one section for treated mice. **(H)** CFU/spleen of each strain in the Δ*ompF* + WT competition experiments. All mouse experiments were carried out on two independent days. Statistics: **(A, B, C, E, F, G)** Mann-Whitney; **(D, H)** Kruskal-Wallis one-way ANOVA with Dunn’s post-test. ****p<.0001, **p<.01, *p<.05, n.s.: not significant.

During co-infection with the WT and Δ*ompF* strains, we confirmed doxycycline treatment significantly decreased CFUs (**Figure 4E**). However, the WT and Δ*ompF* strains were equally fit in the absence and presence of doxycycline, hence while the Δ*ompF* strain may have slight increased susceptibility to doxycycline *in vitro,* this wasn’t sufficient to impact survival of the strain within mouse tissues (**Figure 4F**). The microcolony areas of the WT and Δ*ompF* strains were similar, although the Δ*ompF* microcolonies were slightly larger than WT in untreated mice, but the areas were not significantly altered by treatment (**Figure 4G**). There were few Δ*ompF* microcolonies remaining within host tissues after treatment, which may suggest some increased susceptibility of the strain. However, the number of WT microcolonies in the same spleens also decreased (**Figure 4G**). The CFUs of the WT and Δ*ompF* strain were similar before and after treatment, however the decrease in Δ*ompF* CFUs after treatment was slightly more pronounced, which could indicate a small increase in Δ*ompF* Dox sensitivity *in vivo* (**Figure 4H**).

### Δ*ompF* has altered doxycycline permeability

Increased expression of the OmpF porin in response to Dox was unexpected, since OmpF expression could increase the diffusion of antibiotic into bacterial cells ^36, 39, 40^. We hypothesized that Dox may use a different outer membrane protein for entry into *Y. pseudotuberculosis*, and that increased expression of *ompF* may promote antibiotic diffusion back out of the cell. This would lead to increased levels of Dox accumulation in the Δ*ompF* strain, where Dox could enter the cell, but not diffuse out. Similarly, we hypothesized that OsmB lipoprotein could impact membrane permeability, and sought to determine if there was also differential Dox accumulation in Δ*osmB* cells. However, the limited impact of OsmB on Dox sensitivity meant that any change in Dox accumulation in the Δ*osmB* strain would likely be minimal.

To determine if Dox differentially accumulates in mutant strains, we utilized a TetON reporter system to detect intracellular Dox concentrations. In this reporter system, increasing concentrations of Dox results in heightened levels of mCherry signal, until concentrations are sufficient to inhibit mCherry translation ^17^. This results in a bell curve, or normal distribution, of mCherry expression. The WT and deletion strains (Δ*osmB,* Δ*ompF*, Δ*osmB* Δ*ompF*) were transformed with the TetON reporter plasmid then exposed to increasing concentrations of Dox in culture, and we quantified mCherry fluorescence within individual cells by fluorescence microscopy. In the absence of treatment, background mCherry fluorescence was similar in all strains, but slightly higher in Δ*ompF* (**Figure 5A**). With low concentrations of Dox (0.01µg/ml), we detected high levels of fluorescence in the Δ*ompF* strain, consistent with increased antibiotic accumulation (**Figure 5B**). Interestingly, the Δ*osmB* Δ*ompF* had slightly increased fluorescence compared to the WT strain, but much lower fluorescence than Δ*ompF*, again indicating deletion of *osmB* may have partially rescued the *ompF* phenotype (**Figure 5B**). Consistent with increased antibiotic accumulation in Δ*ompF*, exposure to 0.1µg/ml Dox caused fluorescence to drop below WT levels, suggesting this relatively low concentration was accumulating to sufficient levels to inhibit mCherry translation (**Figure 5C**). Exposure to 1µg/ml Dox resulted in decreased mCherry fluorescence in all mutants relative to WT, suggesting they may all have some level of increased antibiotic accumulation (**Figure 5D**). Throughout these *in vitro* experiments we saw the most dramatic changes with Δ*ompF*, providing evidence this gene product impacts antibiotic accumulation within the cell. The impact of *osmB* was subtle, while the Δ*osmB* Δ*ompF* strain showed an intermediate phenotype; this will be discussed in more detail in the Discussion.

**Figure 5:**
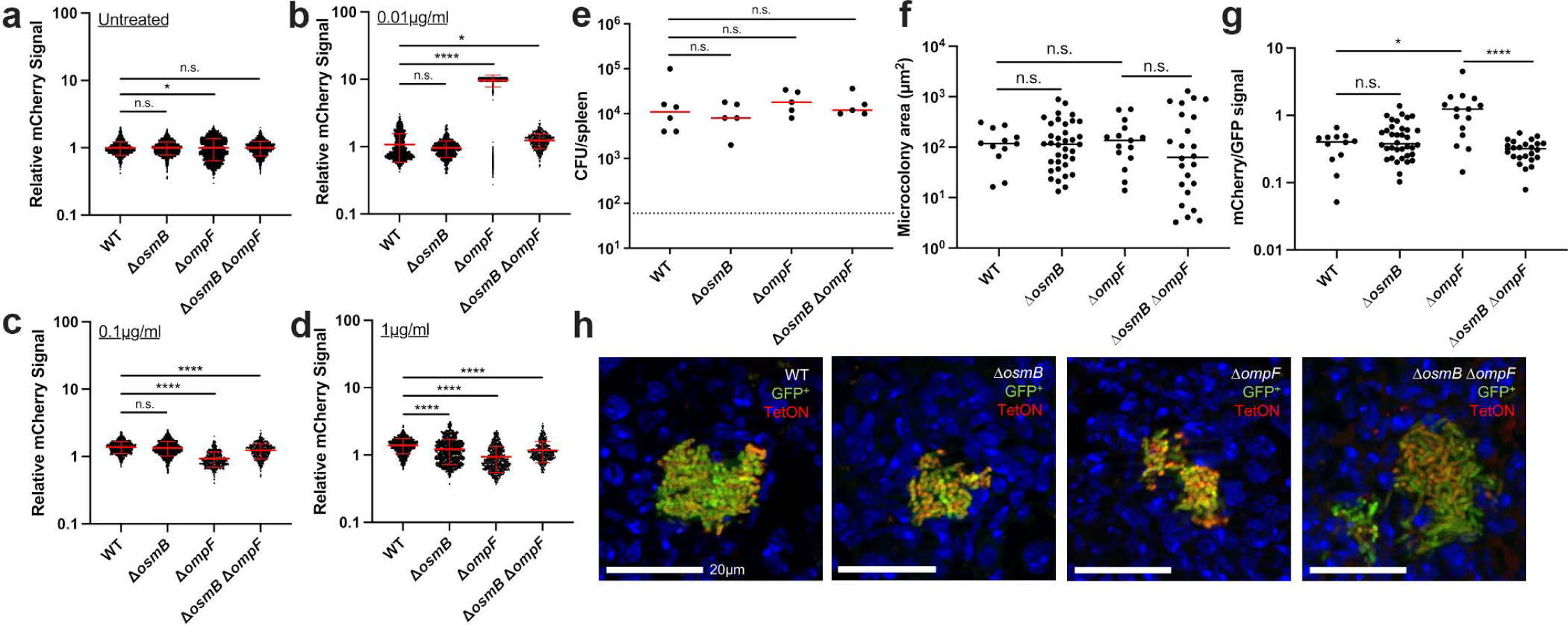
Δ*ompF* has altered doxycycline permeability. **(A-D)** WT and deletion strains (Δ*osmB,* Δ*ompF*, Δ*osmB* Δ*ompF*) transformed with the TetON reporter plasmid were incubated with the indicated concentrations of Dox for 4h and mCherry fluorescence was detected within individual cells by fluorescence microscopy. Relative mCherry was calculated by normalizing to the WT untreated average cell value. Data represents the mean and standard deviation of three biological replicates. **(E)** C57BL/6 mice were infected with WT or mutant strains expressing constitutive GFP alongside the TetON reporter. Infections proceeded 48h, and mice were injected intraperitoneally with a single dose of Dox. Spleens were harvested 24h after treatment to **(E)** quantify CFUs, **(F)** quantify microcolony areas, and **(G)** quantify reporter signal by fluorescence microscopy. Relative reporter signal was quantified by dividing the mean mCherry (TetON) signal by the mean GFP signal from each microcolony. 5 mice were quantified/strain, and 1-15 microcolonies were quantified in one section for each mouse. **(H)** Representative images, scale bars: 20µm. All mouse experiments were carried out on two independent days. Statistics: Kruskal-Wallis one-way ANOVA with Dunn’s post-test. ****p<.0001, *p<.05, n.s.: not significant.

To determine if differential antibiotic accumulation occurs in mutant strains during infection, we constructed GFP^+^ TetON mutant strains. GFP signal will be constitutively expressed, and mCherry will be induced by Dox, which will allow us to assess differential TetON reporter expression across microcolonies ^17^. We performed single strain infections with WT, Δ*osmB,* Δ*ompF*, or Δ*osmB* Δ*ompF* GFP^+^ TetON strains, allowed infection to proceed 48h, injected intraperitoneally with Dox, and harvested spleens 24h after treatment to quantify CFUs, quantify microcolony areas, and detect reporter signal by fluorescence microscopy. The CFUs and microcolony areas for each strain were similar, consistent with the previous findings in Figures 4B and 4E, indicating these strains are not significantly more sensitive to doxycycline than WT (**Figure 5E, 5F**). The relative Dox exposure was then quantified by dividing the mean mCherry signal by the mean GFP signal for each microcolony. Δ*ompF* microcolonies had significantly higher reporter signal relative to the other strains, suggesting that heightened Dox accumulation also occurs within this mutant during infection (**Figure 5G, 5H**). It is important to note that while Δ*ompF* may accumulate more Dox, this is not sufficient to significantly impact bacterial survival (**Figure 5E**) or microcolony area (**Figure 5F**) when comparing across single strain infections, indicating the impact of OmpF on antibiotic susceptibility is subtle. We again observed fewer microcolonies within Δ*ompF* infected tissues; it required sectioning deeper into tissues to find bacterial centers, suggesting the mutant strain may not retain as many microcolonies after treatment.

### *tusB* overexpression promotes *Y. pseudotuberculosis* doxycycline sensitivity *in vitro*

tRNA 2-thiouridine synthesizing protein B (*tusB*) is a protein that functions as a heterohexamer with TusC and TusD in the sulfur-relay system, responsible for the post-transcriptional 2-thiolation of the uracil 34 at the wobble position ^33, 34^. This modification is necessary for the identification of purines, promoting proper translation and incorporation of glutamic acid, glutamine and lysine into growing polypeptide chains ^33^. The RNA-seq results showed that *tusB* is significantly downregulated in response to doxycycline, thus, we chose to generate a tet-inducible (TetON) *tusB* overexpression strain (*P_tetA_::tusB*), to specifically induce *tusB* expression in the presence of doxycycline and prevent downregulation ^17^ (**Figure 6A**). To determine if the Dox MIC was altered by increased expression of *tusB,* we performed growth curves with increasing levels of Dox. Overexpression of *tusB* did not alter bacterial growth kinetics or the MIC of the strain, and we observed similar absorbance levels for the WT and *P_tetA_::tusB* strains under untreated and treated conditions (**Figure 6B-E**).

**Figure 6:**
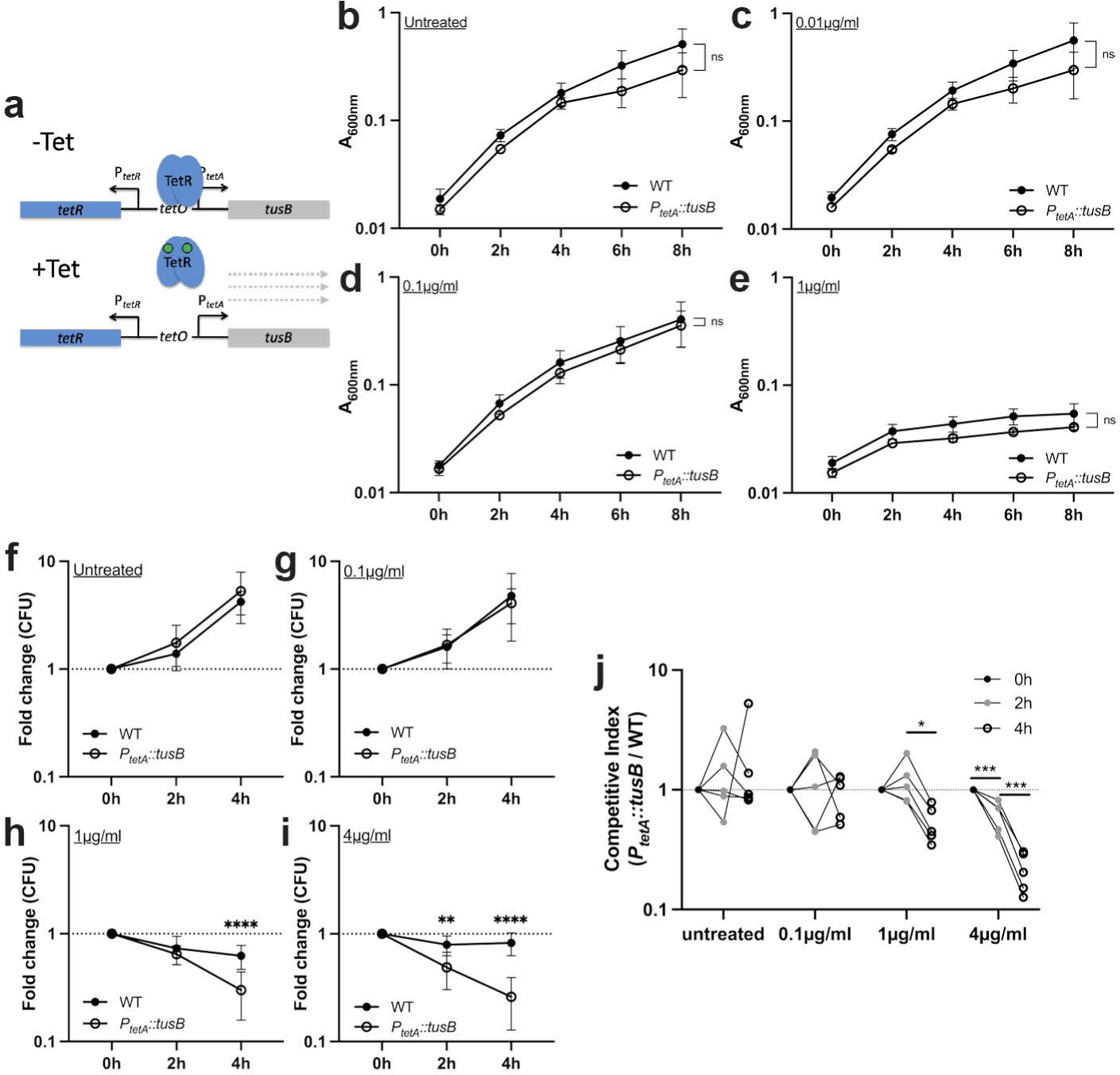
*tusB* overexpression promotes *Y. pseudotuberculosis* doxycycline sensitivity *in vitro.* **(A)** Schematic of *tusB* overexpression strain. *tusB* expression is repressed by TetR in the absence of tetracyclines (-Tet). In the presence of antibiotics (+Tet), TetR repression is relieved, and *tusB* is expressed. Exponential phase cultures of the WT and *tusB* overexpression strain (*P_tetA_::tusB*) were treated with the indicated concentrations of Dox to assess **(B-E)** growth inhibition, **(F-I)** changes in viable counts, and **(J)** competitive survival. **(B-E)** Strains were incubated with the indicated concentrations of Dox and growth inhibition was assessed based on absorbance (A_600nm_) at the indicated timepoints (time: hours, h). Data represents the mean and standard deviation of five biological replicates. **(F-I)** Strains were incubated with the indicated concentrations of Dox and viable counts were determined by quantifying CFUs. Fold change in CFUs is shown relative to time 0h. Dotted line: a value of 1, no change relative to 0h. Data represents the mean and standard deviation of ten biological replicates. **(J)** Competitive survival in the presence of the indicated concentrations of Dox. Competitive index: CFUs of the *P_tetA_::tusB* /WT divided by the ratio of *P_tetA_::tusB* /WT in the culture at time 0h. Values above 1 indicate the *tusB* strain preferentially survives, values less than 1 indicate the WT preferentially survives. Dotted line: value of 1, equal fitness. Dots: biological replicates, lines connect biological replicates sampled across the timepoints, five biological replicates shown. Statistics: Two-way ANOVA with Bonferroni’s multiple comparison test, **(B-I)**: comparisons made relative to the WT strain; **(J)**: comparisons made between timepoints at the same Dox concentration. ****p<.0001, ***p<.001, **p<.01, *p<.05, n.s.: not significant.

We then tested the viability of the WT and *P_tetA_::tusB* strains during exposure to heightened, inhibitory levels of Dox (0.1-4µg/ml). As described above, we exposed exponential phase cells to Dox for 4h, and plated cells at the point of adding Dox (0h), 2h, and 4h after exposure to quantify the number of viable cells during the exposure. Strains were grown in separate test tubes with rotation, and we calculated the fold change in CFUs relative to the 0h timepoint. As seen with the *osmB* and *ompF* mutants, strains continued to grow under untreated conditions and during exposure to 0.1µg/ml Dox **(Figure 6F, 6G**). However, there was a significant drop in the number of *P_tetA_::tusB* cells at 4h of treatment with 1µg/ml Dox compared to WT cells (**Figure 6H**). This phenotype was more pronounced with 4µg/ml Dox, where there was a significant drop in *P_tetA_::tusB* cells relative to WT after only 2h exposure, and greater loss in viability at 4h post-treatment (**Figure 6I**). These data strongly support a role for *tusB* downregulation in Dox tolerance, and show that Dox can have bactericidal activity against the *P_tetA_::tusB* strain. This defines the *P_tetA_::tusB* strain as susceptible, and the WT strain as tolerant under these conditions.

We also tested the competitive survival of the *P_tetA_::tusB* strain relative to WT during co-culture. The *P_tetA_::tusB* construct contains a chloramphenicol resistance cassette, which was used to quantify the CFUs of this strain. An unmarked WT strain was used alongside *P_tetA_::tusB*.

Exponential phase cells were added to individual wells of a 96 well plate, and competitive index values were calculated based on the ratio of *P_tetA_::tusB* /WT CFUs at the indicated timepoints after treatment relative to the ratio when doxycycline was added. These data were consistent with the viability count data, and showed the *P_tetA_::tusB* strain was significantly outcompeted by WT during Dox exposure (**Figure 6J**). The *P_tetA_::tusB* strain lost significantly more viable cells than WT during 4h exposure to 1µg/ml Dox, and at both 2h and 4h after exposure to 4µg/ml Dox. Both methods of detecting antibiotic tolerance showed that *tusB* downregulation significantly contributes to antibiotic tolerance, and that reversing this phenotype by expressing *tusB* during Dox treatment results in heightened drug susceptibility. Bacterial killing was observed in the *P_tetA_::tusB* strain, although we would need to extend experiments longer to accurately determine MBC and MDK values with 90% killing or more stringent cut-offs. However, based on these experiments, approximately 75% of cells were killed with 4µg/ml Dox at 4h, which would mean the MBC75 is ∼4µg/ml and the MDK75 would be ∼4h.

### *tusB* expressing cells have increased susceptibility to doxycycline *in vivo*

To determine if the increased Dox sensitivity of the *P_tetA_::tusB* strain resulted in impaired survival during Dox treatment in the mouse model of infection, we compared the survival of the WT and *P_tetA_::tusB* strains during co-infection. Equal amounts of the WT and *P_tetA_::tusB* strain were mixed, mice were inoculated intravenously, then at 48h post-inoculation mice either received a single dose of doxycycline or were left untreated. At 24h post-treatment, spleens were harvested to quantify CFUs. Enumeration of total CFUs indicated that the Dox was sufficient to significantly decrease CFUs (**Figure 7A**), however the competitive index values were similar with and without treatment (**Figure 7B**). Notably, the competitive index values were below one, indicating the *P_tetA_::tusB* strain was slightly outcompeted by WT in mouse tissues even under untreated conditions, despite very similar fitness in culture. We then quantified the CFUs of each strain before and after treatment within the same tissues. We saw a pronounced reduction in the CFUs of the *P_tetA_::tusB* strain after treatment and also a reduction in WT CFUs, although this was not statistically significant (p = 0.0502) (**Figure 7C**). While the WT and *P_tetA_::tusB* CFUs were similar prior to treatment, there was a significant drop in *P_tetA_::tusB* CFUs relative to WT +Dox treatment, indicating the *P_tetA_::tusB* strain does have heightened sensitivity to Dox within mouse tissues (**Figure 7C**). To determine if this heightened sensitivity also results in smaller microcolony areas after treatment, we transformed WT *Y. pseudotuberculosis* containing *yopE::mCherry* with the *P_tetA_::tusB* plasmid and repeated infections with this strain. Mice were inoculated intravenously with the *yopE::mCherry P_tetA_::tusB* strain, at 48h post-inoculation mice received a single dose of Dox or were left untreated, and spleens were harvested 24h post-treatment. The CFUs of the *yopE::mCherry P_tetA_::tusB* strain significantly decreased with treatment (**Figure 7D**), however this was not reflected by a change in microcolony size (**Figure 7E**). We did observe there were fewer total microcolonies in treated tissues, as we had also seen with the WT, Δ*osmB*, and Δ*ompF* infections. These results could suggest that larger and smaller microcolonies are equally susceptible to treatment, and may be eliminated or survive at a similar rate.

**Figure 7:**
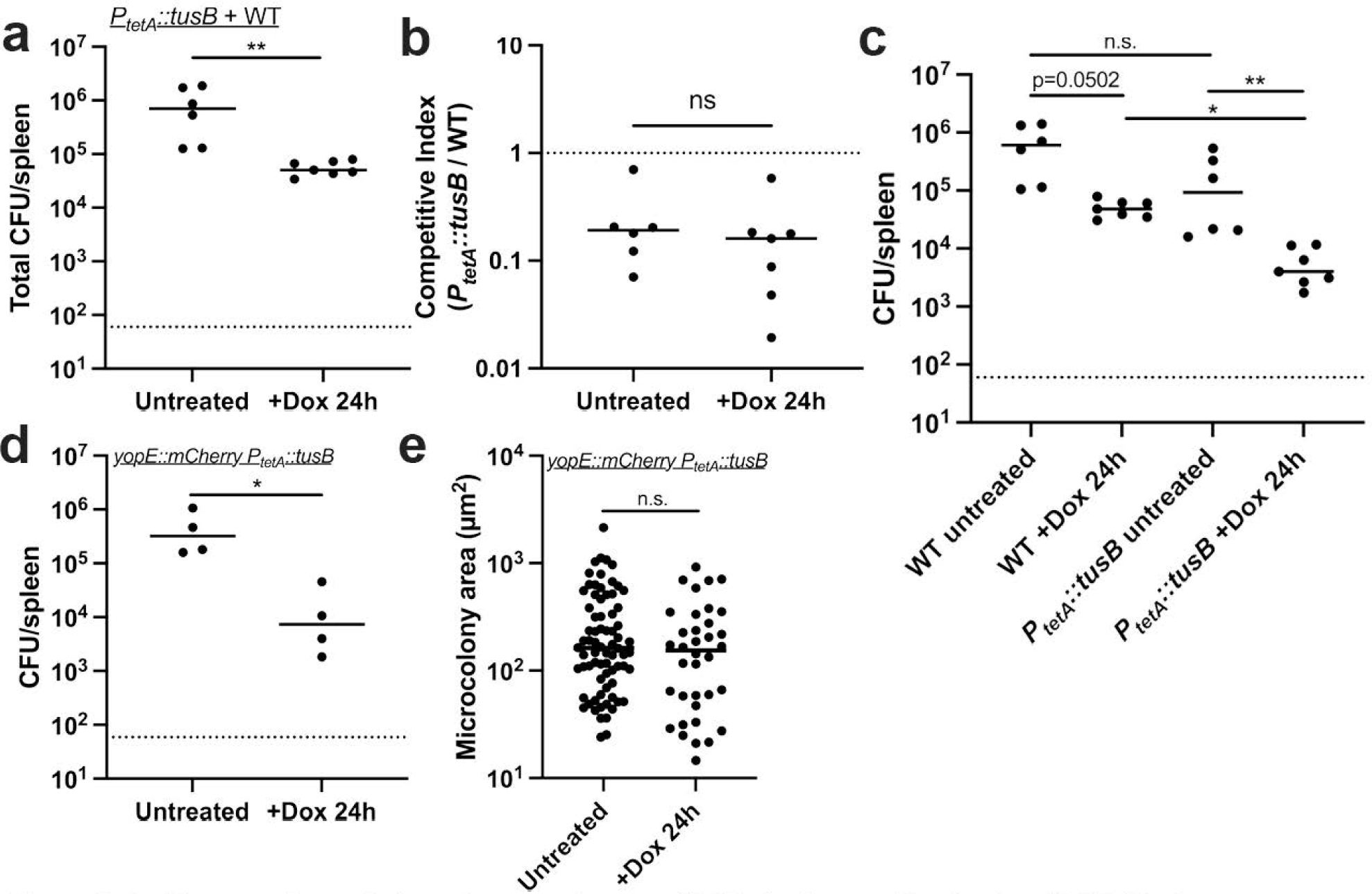
*tusB* expressing cells have increased susceptibility to doxycycline *in vivo.* C57BL/6 mice were infected with equal numbers of the WT and *P_tetA_::tusB* strains, infection proceeded for 48h, and mice were either injected intraperitoneally with a single dose of Dox or left untreated. 24h later spleens were harvested to quantify CFUs. **(A)** Total CFU/spleen, each dot represents an individual mouse. **(B)** Competitive index values. Dotted line: value of 1; equal fitness between the two strains. Values less than 1 indicate the *tusB* strain is less fit than WT. **(C)** CFU/spleen of each strain in the WT + *P_tetA_::tusB* competition experiments. **(D)** C57BL/6 mice were infected with the *yopE::mCherry P_tetA_::tusB* strain, infection proceeded for 48h, and mice were either injected intraperitoneally with a single dose of Dox or left untreated. 24h later spleens were harvested to quantify CFUs. **(E)** Microcolony areas of the *yopE::mCherry P_tetA_::tusB* strain were quantified from 4 untreated mice and 4 treated mice, each dot represents an individual microcolony. 6-38 microcolonies were quantified in one section for untreated mice, and 1-19 microcolonies were quantified in one section for treated mice. All mouse experiments were carried out on two independent days, horizonal lines depict median values. Statistics: **(A, B, D, E)** Mann-Whitney; **(C)** Kruskal-Wallis one-way ANOVA with Dunn’s post-test. **p<.01, *p<.05, n.s.: not significant.

### *ompF, tusB,* and *cnfy* are differentially regulated in response to inhibitory concentrations of chloramphenicol, while *osmB* expression was only altered by doxycycline

Many antibiotics target the ribosome to inhibit translation and arrest bacterial growth. To begin to determine if the differential regulation of *osmB, ompF, tusB* and *cnfy* was a response to doxycycline, or a more general response to ribosomal inhibition, we also performed experiments with chloramphenicol (Cm), which targets the 50S ribosomal subunit by binding to 23S rRNA ^53^. Cm was also chosen for these experiments because it can be used to treat *Yersinia* infections ^20, 22^. We performed growth curves with increasing doses of Cm (0.0025µg/ml-50µg/ml) to determine the sensitivity of WT *Y. pseudotuberculosis* to Cm, and define the minimum inhibitory concentration (MIC). All concentrations tested resulted in significant growth inhibition after 6 hours of growth in bacteriological media (**Figure 8A**). However, the growth curve with 0.0025µg/ml chloramphenicol mirrored untreated growth through 4 hours of growth, suggesting this low dose minimally impacted growth, and was sub-inhibitory. Treatment with 0.025-0.25µg/ml resulted in similar levels of growth inhibition, and treatment with 2.5-50µg/ml resulted in severe growth inhibition, to the point that no growth was detected (**Figure 8A**). Based on these data, the MIC of our WT *Y. pseudotuberculosis* strain to Cm was 2.5µg/ml.

**Figure 8:**
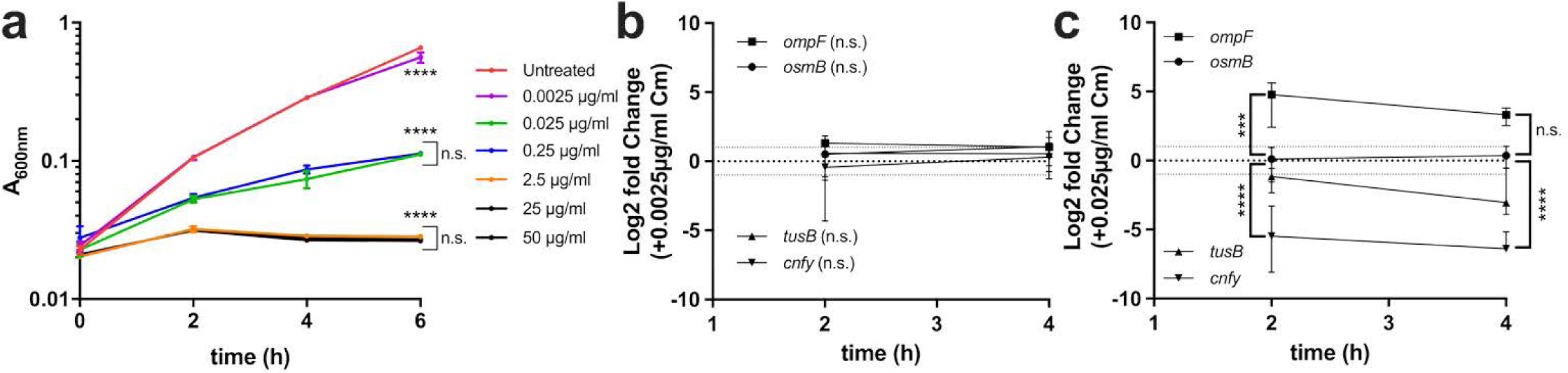
*ompF, tusB,* and *cnfy* are differentially regulated in response to inhibitory concentrations of chloramphenicol, while *osmB* expression was only altered by doxycycline. **(A)** WT *Y. pseudotuberculosis* was grown in the presence of the indicated concentrations of chloramphenicol (Cm). Growth was measured by absorbance (A_600nm_) at the indicated timepoints (hours, h). Bacteria were grown in the presence or absence of **(B)** 0.0025µg/ml Cm or **(C)** 0.025µg/ml Cm. RNA was isolated after 2h and 4h exposure, and the indicated transcripts were detected by qRT-PCR. Fold change is relative to untreated cells (represented by a dotted line at a value of 1). Statistics: **(A-C)** Two-way ANOVA with Tukey’s multiple comparison test, comparisons made relative to untreated cells, comparisons between treatment conditions where indicated by brackets. In C, statistics represent treated vs. untreated comparisons for *cnfy*, and *ompF* compared to *osmB*, which did not change with treatment. ****p<.0001, ***p<.001, n.s.: not significant.

We then compared the levels of *osmB*, *ompF*, *tusB* and *cnfy* transcripts during exposure to the subinhibitory 0.0025µg/ml Cm dose and the inhibitory 0.025µg/ml dose. The subinhibitory 0.0025µg/ml Cm did not significantly change any transcript levels when comparing untreated and treated cells (**Figure 8B**). Interestingly, the inhibitory 0.025µg/ml Cm dose did not change *osmB* transcript levels, which may suggest that the change in *osmB* transcripts in response to Dox is antibiotic-specific, and not a general response to ribosomal inhibition (**Figure 8C**). *ompF* was significantly upregulated in response to this concentration of Cm, and *cnfy* was significantly downregulated, suggesting these could be general responses to ribosomal inhibition. *tusB* transcript levels also decreased in response to Cm, although this was not statistically significant. Collectively, these data suggest that transcriptional regulation of *ompF, cnfy,* and *tusB* may be a response to ribosomal targeting by antibiotics, while heightened *osmB* expression may be a specific response to doxycycline.

## Discussion

Antibiotic tolerance is associated with a phenotypic change within a bacterial population that results in decreased antibiotic susceptibility. Tolerance is specifically associated with bactericidal antibiotics, however, under conditions where bacteriostatic antibiotics are directly killing cells, the terminology can also be applied to a bacteriostatic antibiotic, such as doxycycline (Dox) ^14–16^. Here, we sought to determine how bacteria respond to Dox exposure in culture, and determine whether differential gene expression is required for surviving Dox treatment in culture and in our mouse model of *Y. pseudotuberculosis* infection. Only four genes were differentially regulated ∼2 fold in response to a physiologically-relevant dose of Dox (**Figure 1B**): *osmB, ompF, tusB* and *cnfy;* differential gene expression occurred in response to low and high doses of Dox and during antibiotic treatment in our mouse model (**Figure 2A, 2B**). We then tested the role of *osmB*, *ompF*, and *tusB* differential expression in doxycycline tolerance by generating deletion mutants for the genes with heightened expression (*osmB*, *ompF*), and a *P_tetA_::tusB* construct to specifically promote heightened *tusB* expression during Dox exposure. Using these strains, we found that *osmB* expression does not significantly contribute to Dox susceptibility, *ompF* expression impacts Dox accumulation within bacterial cells and has a slight contribution to tolerance, while *tusB* downregulation significantly contributes to Dox tolerance. *cnfy* expression has previously been associated with persistence in *Y. pseudotuberculosis* ^28^, and so we chose to focus on *osmB, ompF,* and *tusB* in our experiments here. Interestingly, *ompF, tusB* and *cnfy* were also differentially regulated in response to chloramphenicol, suggesting these could be general responses to ribosomal inhibition, while *osmB* upregulation may be more specific to Dox (**Figure 8C**).

Expression of the *osmB* lipoprotein slightly impaired bacterial growth, based on *in vitro* susceptibility experiments and the improved fitness of the Δ*osmB* strain in the mouse model, independent of drug treatment (**Figure 4B, 4C**). These data may suggest there is redundancy in lipoprotein expression, and that expression of other lipoproteins are sufficient to compensate for a loss of *osmB. osmB* expression increases during a transition to stationary phase growth, and in response to changes in osmolarity, which would presumably promote increased membrane fluidity under these conditions ^43, 44, 54^. Multiple *osm* genes are expressed under these conditions, and subsets of these could potentially compensate for loss of *osmB* ^41^. OsmB also has a role in repairing the *E. coli* outer membrane after stress, which would suggest that OsmB could play some role in membrane repair after damage caused by antibiotic treatment ^43, 54, 55^. Although our data suggest that OsmB does not significantly contribute to Dox susceptibility or tolerance in the conditions tested, as loss of *osmB* only had a slight impact on the accumulation of Dox within bacterial cells and this did not alter the susceptibility of the Δ*osmB* strain, OsmB may still play some other role that helps the cell persist during antibiotic exposure.

*ompF* porin expression was heightened during Dox exposure, which impacted antibiotic accumulation within the bacterial cell, and indicated that OmpF may promote diffusion of doxycycline out of the cell (**Figure 5B, 5G**). These results were surprising; we had expected to see decreased expression of porins during Dox exposure, since OmpF has been shown to promote passive uptake of multiple different antibiotics through the outer membrane, including tetracyclines ^23, 36, 38, 40^. Increased OmpF expression would have been expected to increase the amount of Dox influx into the cell, and negatively impact bacterial survival. The accumulation of Dox in the *ompF* deletion mutant instead indicates this porin may also have a role in passive diffusion of tetracyclines back out of the bacterial cell. Notably, the phenotypes with Δ*ompF* were most pronounced at low levels of Dox, where you might expect more of a role of passive diffusion in lowering intracellular antibiotic levels ^36, 38^. Expression of efflux pumps promotes tetracycline resistance by pumping antibiotics out of the bacterial cell ^23, 25, 36^, and we had also expected to see heightened expression of efflux pumps in response to Dox exposure. It is possible that we might see expression of efflux pumps in response to heightened levels of Dox, and it is likely that many additional pathways are differentially regulated in response to more stressful, inhibitory concentrations of Dox. It will be interesting to further characterize the transcriptional changes that occur during exposure to heightened doses of Dox, and also probe more deeply the bacterial responses to Dox treatment within host tissues.

Another surprising result was the comparison in phenotypes of the Δ*ompF* and Δ*osmB* Δ*ompF* strains. After seeing subtle phenotypes with the Δ*ompF* strain, we had expected to see similar or stronger evidence of antibiotic susceptibility with the Δ*osmB* Δ*ompF* strain. Instead, the deletion of *osmB* appeared to partially rescue the Δ*ompF* phenotype, possibly indicating aberrant *osmB* expression, detrimental mislocalization of OsmB, or misregulation of other outer membrane components, occurs in the absence of OmpF ^56^. A direct interaction between OmpF and OsmB has not been shown, however it is possible OsmB could function like other lipoproteins, and contribute to phospholipid transport through interactions with porins ^56^. In which case, loss of OmpF could have additional effects on the outer membrane, which could explain a stronger phenotype with the Δ*ompF* single deletion strain. It is also interesting to think about the expression of these two genes, and what it means to see both genes upregulated in response to Dox. Both *osmB* and *ompF* are regulated by the alternative sigma factor, RpoS, which promotes expression of genes in stationary phase ^40, 54^. However, *osmB* expression occurs independently of EnvZ and OmpR, which are required for *ompF* expression, again suggesting *osmB* and *ompF* are regulated independently ^41^. It has also been shown that *osmB* is expressed under high osmolarity, conditions that repress *ompF* expression, providing further evidence that expression of these genes does not typically occur under the same conditions ^41^. These data indicate that Dox exposure does not mimic different changes in osmolarity, and while it is possible that RpoS-dependent gene expression could occur in response to this stress, we did not see heightened expression of the RpoS-regulated gene, *dps* (**Supplemental Figure 1**).

Overexpression of *tusB* resulted in a significant impact on bacterial viability (**Figure 6H, 6I, 6J, 7C**), suggesting that tRNA modification by *tusB*, and resulting impacts on translational machinery, may play an important role in promoting tolerance. We believe this may be the first time bactericidal activity of Dox has been described, however it is possible that other genetic modifications could also promote bactericidal activity of this typically bacteriostatic antibiotic ^14, 15^. Tetracyclines bind reversibly to the 16S rRNA component of the ribosomal small subunit, inhibiting translation ^24^. TusB function, in contrast, promotes the stabilization of the wobble position of specific tRNAs for proper incorporation of glutamine (Gln), glutamate (Glu), and lysine (Lys) residues into growing polypeptide chains ^33^. Because of the role of TusB relatively early in this 2-thiouridylation cascade, it is unlikely that TusB remains associated with tRNAs as they enter ribosomes, and so Dox and TusB should not both be interacting with ribosomes directly ^33, 34^. Instead, we believe that the toxicity associated with heightened TusB expression during Dox exposure could be linked to heightened activity of the cascade, increased levels of 2-thiouridylation, and potential stalling at Gln, Glu, and Lys codons for proper incorporation of these amino acids. This could result in slower translation of proteins, and a slower response to Dox exposure. Conversely, in WT cells that downregulate *tusB* expression, we would expect cells to translate proteins more quickly, albeit with errors in incorporation of Gln, Glu, and Lys residues. We are very interested in pursuing the mechanism underlying the TusB phenotype in future studies, to better understand how modulation of translation may promote increased Dox toxicity, and ultimately improved efficacy of Dox treatment. Based on previous studies, the observed bactericidal activity is likely due to prolonged Dox binding to ribosomes resulting in fewer active ribosomes ^14, 15^. It is also important to note there was a slight fitness cost associated with the *P_tetA_::tusB* construct *in vivo* that may be due to increased *tusB* expression within mouse tissues prior to Dox administration. This was specific to the host environment, since the *P_tetA_::tusB* strain was as fit as the WT strain during growth in culture. For future studies we plan to generate a panel of different *tusB* strains to examine the impact of constitutive *tusB* expression, induction from a single copy *P_tetA_::tusB* construct, and deletion of *tusB* on Dox susceptibility.

We have found in previous studies that bacteria within microcolonies experience different microenvironments, resulting in differing levels of stress responses ^51, 57^. We have also shown that Dox does not diffuse evenly across microcolonies at this 40mg/kg single injection dosage ^17^, which suggests that bacteria may also respond to Dox in a heterogeneous manner within host tissues. It remains unclear if the genes we have studied here are differentially regulated within the same cells, or if distinct subpopulations differentially express these gene products. In future experiments, we plan to explore this idea using fluorescent reporters to determine whether individual cells differentially regulate genes of interest in response to Dox, and also develop single cell assays to better understand and characterize the extent of heterogeneity in the response to antibiotics. It will also be important to characterize the response to Dox specifically within host tissues. For additional future studies, we plan to isolate surviving bacterial cells from the host environment to determine which pathways promote persistence in the host, and compare these to the specific responses to Dox in culture.

## Materials and Methods

### Bacterial strains & growth conditions

The WT *Y. pseudotuberculosis* strain, IP2666, was used throughout this study. For all *in vitro* experiments, overnight cultures of bacteria were grown for 16-18 h in lysogeny broth (LB) at 26°C with rotation. Bacteria were then sub-cultured 1:100 into fresh LB and incubated at 37°C with rotation for experiments. Doxycycline was added to the samples at the indicated concentrations. For *in vivo* experiments, bacteria were cultured overnight as described above, in 2xYT broth (LB with 2x yeast extract and tryptone) ^17, 51^. Bacteria were then diluted in sterile phosphate-buffered saline solution (PBS) to a final concentration of 10^3^ CFU/100µl.

### qRT-PCR to detect bacterial transcripts in broth-grown cultures

Bacterial cells were grown in the presence or absence of antibiotics for the indicated time points, pelleted, resuspended in Buffer RLT (QIAGEN) + ß-mercaptoethanol, and RNA was isolated using the RNeasy kit (QIAGEN). DNA contamination was eliminated using either the Turbo DNA-free kit (Invitrogen) or the on-column RNase-free DNase set (QIAGEN). RNA was reverse transcribed using M-MLV reverse transcriptase and random primers (Invitrogen), in the presence of the RNase inhibitor, RnaseOUT (Invitrogen). Approximately 30 ng cDNA was used as a template in qPCR reactions with 0.5 µM of forward and reverse primers ^51^ and Power SYBR Green PCR Master mix (Applied Biosystems). Control samples were prepared that lacked reverse transcriptase to confirm genomic DNA was eliminated from samples. qPCR reactions were carried out using the StepOnePlus Real-Time PCR system, and comparisons were obtained using the ΔΔCT and 2^-ΔCt^ method (Applied Biosystems) for relative expression. Kits were used according to manufacturers’ protocols.

### RNA-seq

Overnight cultures of WT *Y. pseudotuberculosis* were diluted 1:100 into fresh media and grown at 37°C in the presence or absence of 0.1µg/ml Dox. At 2h and 4h post-treatment, approximately 10^5^ bacterial cells were collected from cultures, and RNA was isolated as described above. RNA was prepared for RNA-seq analysis using a RNAtag-seq approach ^35^. Following sequencing, reads were aligned using the *Y. pseudotuberculosis* strain YPIII fully annotated genome (genome: NC_010465, virulence plasmid: NZ_LT596222). Relative abundance changes in transcript levels were assessed using a DESeq2 method of pairwise comparisons between treatment groups ^58^. An adjusted p-value of 0.05 was considered significant, and we focused on genes differentially regulated ≥2 fold for validation experiments and downstream assays.

### Murine model of systemic infection

Jackson Laboratories (Bar Harbor, ME) provided 6–8 week-old female C57BL/6 mice and the animal experiments were approved by the Johns Hopkins University Institutional Animal Care and Use Committee. Mice were inoculated intravenously via tail vein injection with 10^3^ CFU *Y. pseudotuberculosis* in 100µl for all experiments ^51^. At 48 hours post-inoculation, mice were treated with 40 mg/kg doxycycline (720µg in 100µl sterile PBS) via intraperitoneal injection ^17^. Spleens were harvested at the indicated timepoints post-treatment and either homogenized to quantify CFUs, stabilized in RNALater for RNA isolation, or fixed in 4% paraformaldehyde (PFA) for histology. Tissue was fixed by incubating 24h in 4% PFA at 4° C.

### qRT-PCR to detect bacterial transcripts from infected tissues

At the indicated timepoints, spleens were harvested and immediately submerged in RNALater to stabilize transcripts, incubated at 4°C three weeks, then tissue was stored at –80°C prior to RNA isolation. Tissue was then thawed, added to Buffer RLT (QIAGEN) + ß-mercaptoethanol (200µl/10mg tissue), and homogenized. Cell debris was pelleted, and lysate was used for RNA isolation as described above. DNA contamination was eliminated using the Turbo DNA-free kit (Invitrogen). The MicrobEnrich kit (Invitrogen) was used to deplete mouse mRNAs, tRNAs and rRNAs. Approximately 500ng RNA was used in reverse transcription reactions, and approximately 25ng cDNA were used in qPCR reactions, as described above.

### Generation of deletion strains

*osmB* and *ompF* deletion strains were generated by amplifying the start codon + 3 downstream codons, the 3’ terminal 3 codons + the stop codon and fusing these fragments to generate a start + 6 aa + stop deletion construct for each gene. Deletion constructs were amplified with 500 base pairs flanking sequence on each side, cloned into the suicide vector, pSR47S, and conjugated into *Y. pseudotuberculosis* ^59^. Sucrose selection was used to select for bacteria that had incorporated the desired mutation after a second recombination event ^59^. PCR, sequencing, and qRT-PCR were used to confirm deletion strains. To mark mutant cells with fluorescence, we also generated deletion strains that contained the previously described, *yopE::mCherry* reporter ^51, 52^.

### Generation of the TetON reporter strains and P_tetA_::tusB construct

The TetON reporter (*P_tetA_::mCherry-ssrA*) was previously described ^17^, and was constructed by fusing *tetR* through the *P_tetA_* promoter to a ssrA-tagged destabilized variant of *mCherry* ^60, 61^. The TetON reporter was expressed from the low copy pMMB67EH plasmid and transformed into WT and mutant strains by electroporation ^59^. The constitutive GFP plasmid used alongside TetON in mouse infections has been previously described ^17, 51, 52^. This GFP plasmid was also used to mark the WT strain for *in vitro* and *in vivo* competition experiments with the *osmB* and *ompF* deletion strains. *P_tetA_::tusB* was generated the same way as the TetON reporter; the *tetR* through *P_tetA_* fragment was fused to *tusB*. *P_tetA_::tusB* was inserted into the low copy pACYC184 plasmid and electroporated into WT *Y. pseudotuberculosis*, as previously described ^59^. To mark the *P_tetA_::tusB* strain with fluorescence, we also transformed the *P_tetA_::tusB* plasmid into a WT *yopE::mCherry* reporter strain ^51, 52^.

### Antibiotic susceptibility assays

Antibiotic susceptibility of mutant strains was assessed using three *in vitro* approaches: growth inhibition, viability, and competitive survival. Growth inhibition was assessed by diluting overnight cultures (1:100) into fresh LB containing increasing doses of Dox (0.01-1µg/ml), and quantifying absorbance (600nm) every 2h using a microplate reader. Cultures were incubated for 8h at 37° C with rotation. Viability was assessed with exponential phase cells, which were subcultured from overnight cultures (1:100) into fresh LB and rotated at 26° C for 2h. Exponential phase cells were then transferred to fresh LB containing increasing doses of Dox (0.1-4µg/ml), and incubated 4h at 37° C with rotation. Cells were plated for viability at the time of Dox addition (0h), 2h, and 4h post-treatment. Competitive survival was assessed with exponential phase cells prepared as described above, and cells were co-cultured in wells of 96 well plates with increasing doses of Dox. Plates were incubated at 37° C on an orbital shaker (VWR). Single strain controls were grown in parallel to ensure growth and Dox sensitivity was similar in 96 well plates compared to test tubes. Cells were plated to quantify CFUs of each strain at the time of Dox addition (0h), 2h, and 4h post-treatment. Chloramphenicol (Cm) resistance was used to quantify the number of WT cells relative to total bacterial numbers for the experiments with deletion strains. For the WT vs. *P_tetA_::tusB* experiments, the *tusB* strain was marked with Cm resistance.

### Fluorescence microscopy – host tissues

After fixation with PFA, spleens were frozen-embedded in optimal cutting temperature (O.C.T.) compound (Tissue-Tek, VWR). 10 µm thick sections were obtained with a cryostat microtome (Microm HM 505e) and transferred to charged microscope slides (VWR). Slides were stored at –80° C, then thawed in PBS at room temperature, washed 3x in PBS, and stained with Hoechst (1:10,000 dilution in PBS) for 5 minutes. Slides were then washed 3x with PBS and coverslips were mounted with ProLong Gold (Life Technologies). One section was imaged for each tissue. Tissues were imaged with a Zeiss Axio Observer 7 (Zeiss) inverted fluorescent microscope with an Apotome.2 (Zeiss) with the 63x oil immersion objective. Images of the sections were captured with an Axiocam 702 mono camera (Zeiss).

### Antibiotic permeability assays

Exponential phase cultures of TetON-containing strains (WT, Δ*osmB,* Δ*ompF*, Δ*osmB ΔompF*) were incubated in the absence or presence of increasing doses of Dox (0.01-1µg/ml) for 4h at 37° C with rotation. Approximately 10^6^ cells were pelleted and fixed in 4% PFA at 4° C for 18h (overnight). Fixed cells were pelleted, PFA was removed, and cells were resuspended in 20µl sterile PBS. 5µl of each sample was imaged on 1% agarose (in PBS) gel pads. 5 fields of view were captured for each sample, and single cell fluorescence was quantified using Volocity software, as described below. This resulted in approximately 500-800 total cells imaged across three biological replicates, or approximately 150-300 cells per condition, per replicate.

### Image analysis

Volocity image analysis software was used to quantify microcolony areas as previously described ^17, 51^. Briefly, either GFP or mCherry signal were used to detect individual bacteria and quantify total area of each microcolony. One cross-section/tissue was imaged to represent each spleen. Dox permeability was detected using TetON reporter expression, which was quantified using Volocity to measure single cell mCherry fluorescence. Relative mCherry signal was calculated by dividing the mCherry fluorescence value of each cell by the average mCherry value of untreated WT cells, which were used to set a baseline fluorescence value. For *in vivo* experiments, relative Dox accumulation was quantified by dividing the mean mCherry signal across a microcolony, by the mean signal of constitutive GFP, as previously described ^17^.

## Author Contributions

Conceptualization: HSAM, RKD, JV, KMD; Formal Analysis: HSAM, RKD, JV, KC, BN, ZZ, JA, TV, KMD; Funding Acquisition and Supervision: KMD; Investigation: HSAM, RKD, JV, KC, BN; Methodology: HSAM, RKD, JV, KC; Writing – Original Draft Preparation: HSAM, RKD, JV, KC, KMD; Writing – Review & Editing: HSAM, RKD, JV, KC, BN, ZZ, JA, TV, KMD.

## Acknowledgments

We thank the members of the Davis lab for constructive feedback and suggestions during manuscript preparation. We thank Ralph Isberg for invaluable feedback throughout. This work was supported by a NIAID K22 Career Transition Award (K22 AI123465) and NIAID R21 AI154116 grants to KMD.

---

**Supplemental Figure 1:**
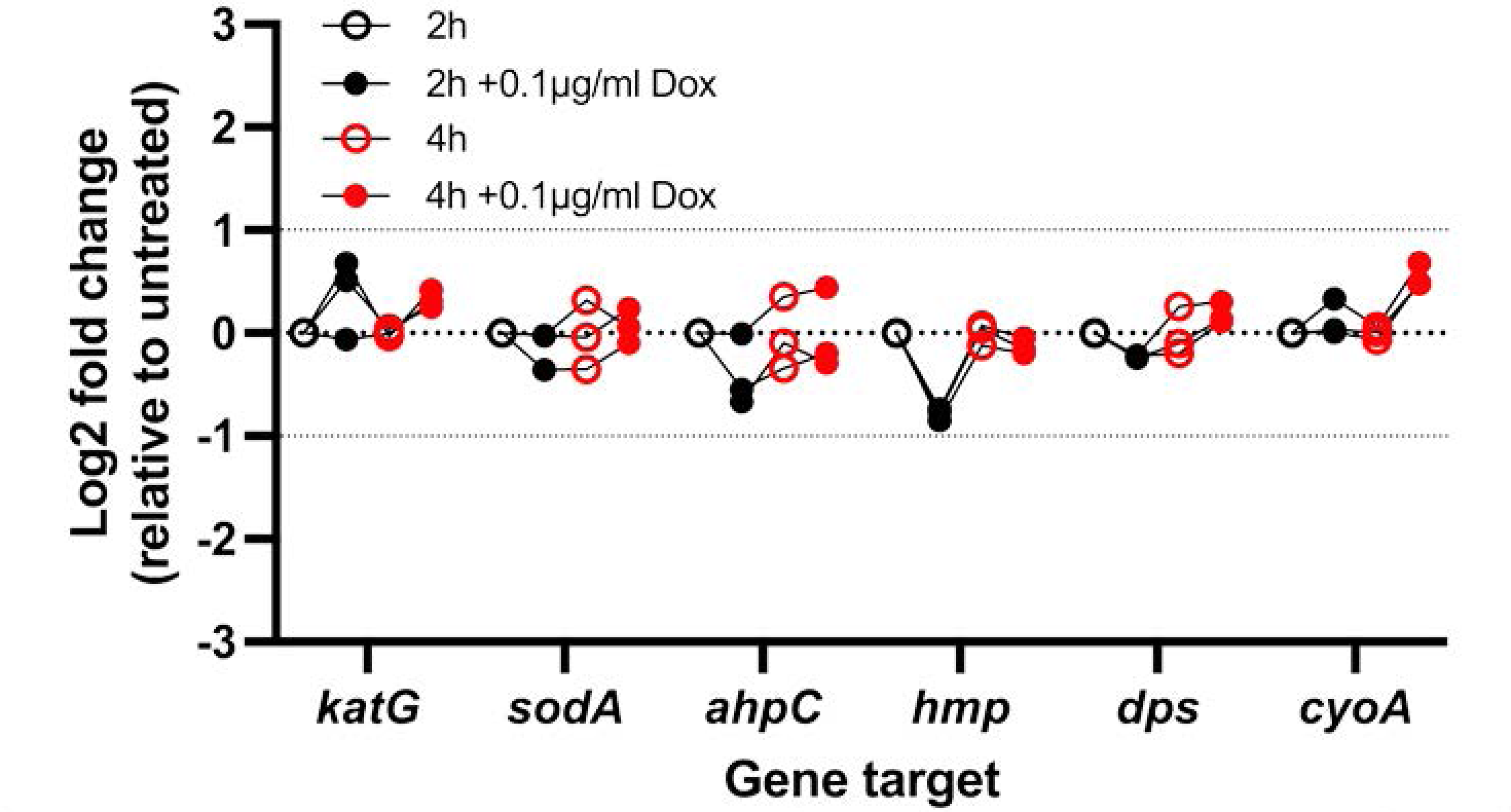
Known stress response genes are not differentially regulated in response to doxycycline. Cultures of WT *Y. pseudotuberculosis* were incubated in the presence or absence of 0.1µg/ml Dox, RNA was isolated and transcripts were detected by qRT-PCR. Log2 fold change values were calculated relative to untreated cells, horizontal dotted lines depict average values for untreated cells (0log2), and 2-fold changes. Data represents four biological replicates.

